# Ventral hippocampal CCK interneurons gate context-reward memory

**DOI:** 10.1101/2023.04.27.538595

**Authors:** Robin Nguyen, Sanghavy Sivakumaran, Evelyn K. Lambe, Jun chul Kim

## Abstract

Associating contexts with rewards depends on hippocampal circuits, with local inhibitory interneurons positioned to play an important role in shaping activity. Here, we hypothesize that the encoding of context-reward memory requires a ventral hippocampus (vHPC) to nucleus accumbens (NAc) circuit that is gated by CCK interneurons. In a sucrose conditioned place preference (CPP) task, optogenetically inhibiting vHPC-NAc terminals impaired the acquisition of place preference. Transsynaptic rabies tracing revealed vHPC-NAc neurons were monosynaptically innervated by CCK interneurons. Using intersectional genetic targeting of CCK interneurons, *ex vivo* optogenetic activation of CCK interneurons increased GABAergic transmission onto vHPC-NAc neurons, while *in vivo* optogenetic inhibition of CCK interneurons increased cFos in these neurons. Notably, CCK interneuron inhibition during sucrose CPP learning increased time spent in the sucrose-associated location, suggesting enhanced place-reward memory. Our findings reveal a previously unknown hippocampal microcircuit crucial for modulating the strength of contextual reward learning.

## INTRODUCTION

Animal survival depends on accurate internal representations of rewarding environments. These contextual memories emerge from the hippocampus^1,2^, yet the neuronal mechanisms underlying their formation remain largely undefined. In particular, the ventral hippocampus (vHPC) is thought to be recruited to encode spatial contexts where emotionally salient events have occurred^3-5^. The spatial firing fields of neurons in the vHPC are typically broad^6,7^, effectively mapping entire contexts^8–10^. Additionally, vHPC neurons display selective firing patterns in situations that are associated with emotional valence and that elicit affect-driven behaviors, including behaviors associated with rewards^8–12^, and avoidance and defensive behaviours^4,8,13–15^. Although there have been several studies that show how neuronal activity is related to reward-associated contexts in the ventral hippocampus (vHPC), the specific circuit mechanisms that are responsible for contextual memory of rewards remain poorly understood.

The activation of neurons in the vHPC region varies across different emotional contexts. Specifically, the involvement of pyramidal neurons may be influenced by their projection targets^10,16^, which are highly heterogeneous^10,17,18^. These projection-defined pyramidal neurons in the vHPC may represent distinct functional populations that are recruited based on task demands^10,13^. Notably, outputs of the vHPC to the nucleus accumbens (NAc) have been found to mediate reward-related goal-directed behaviours^10,18–22^. However, the upstream selection factors allowing for their recruitment during behaviour have not been clearly elucidated.

In addition to the heterogeneity among pyramidal neurons, the hippocampus contains a variety of GABAergic interneurons^23^ that are classified into distinct cell types based on their unique combination of molecular markers, electrophysiological properties, morphology, and connectivity^24–26^. It has been suggested that the varied features among these cell types enable circuit and behavioral specialization^27–33^. Specific classes of GABAergic interneurons may interact with functional groups of hippocampal pyramidal cells to facilitate finely tuned spatial representations underlying memory^30,34–36^.

We have previously shown that vHPC GABAergic interneurons expressing the pan-GABA marker GAD65 are comprised largely of cholecystokinin (CCK) interneurons and play a critical role in regulating dopamine-dependent behaviours^27^. CCK interneurons are highly abundant throughout the septotemporal axis of the hippocampus^37^, and studies have suggested their relevance for spatial memory^38–41^. Therefore, we hypothesize that the vHPC microcircuit involving CCK interneurons and nucleus accumbens-projecting pyramidal neurons (vHPC-NAc) contributes to the formation of contextual reward memory. We used a combination of dual-recombinase intersectional genetics, rabies-mediated monosynaptic tracing, ex vivo electrophysiological recordings, and optogenetic inhibition to target, record, and manipulate the activity of CCK interneurons and vHPC-NAc pyramidal neurons. Our findings demonstrate that local CCK interneurons directly inhibit vHPC-NAc neurons and regulate learning of context-reward associations.

## RESULTS

### vHPC-NAc circuit is required for contextual reward memory

The vHPC may participate in contextual reward memory by associating specific spatial contexts with rewarding stimuli. This function has been attributed particularly to pyramidal neurons which project to the NAc^10,21^. To test whether the vHPC-NAc circuit is involved in the formation of contextual reward memory, we optogenetically inhibited vHPC terminals in the NAc as mice performed a sucrose conditioned place preference task (CPP). AAV containing archaerhodopsin (ArchT) or a reporter (EYFP) driven by the CaMKIIα promoter (AAV2/5-CaMKIIα-eArchT3.0-EYFP or AAV2/5-CAMKIIα-EYFP) was infused bilaterally into the vHPC and optic fibres were implanted above the NAc shell where the densest vHPC projections were observed (Fig. 1A).

**Figure 1.**
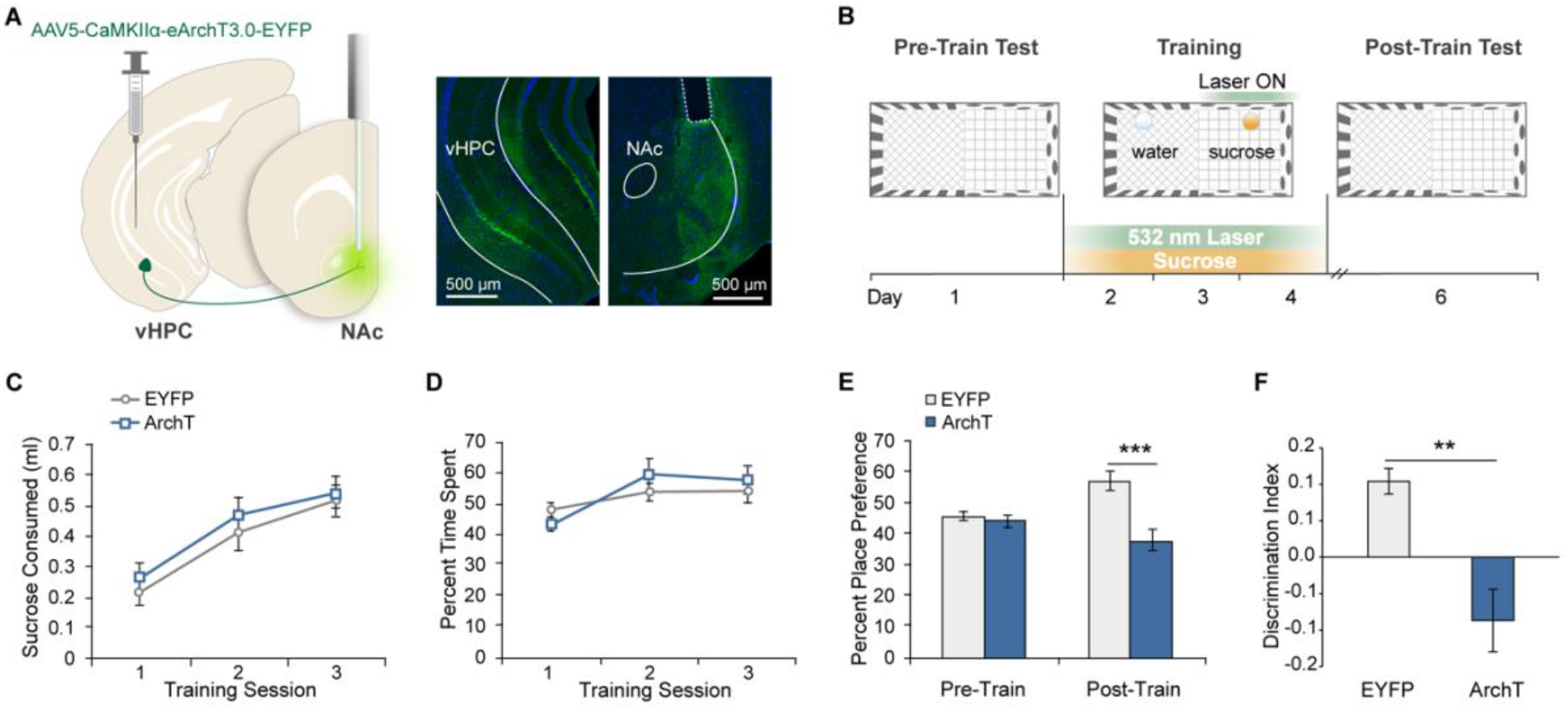
Optogenetic inhibition of vHPC terminals in the NAc during sucrose CPP training impairs memory at test. ***A***. Left, schematic of vHPC-NAc optogenetic circuit targeting strategy. Right, coronal slices of vHPC and NAc expressing eArchT3.0-EYFP (green). ***B***. Behavioral paradigm for closed-loop optogenetic light delivery during sucrose conditioned place preference. Light and sucrose were presented during training sessions only. ***C***. Volume of sucrose solution consumed across training sessions. Mixed ANOVA, no main effect of group *F*(1, 15) = 0.501, *P =* 0.486, no group × day interaction, *F*(2, 30) = 0.24, *P =* 0.786. ***D***. Percentage of time spent in the light/sucrose-paired context across training sessions. No main effect of group, *F*(1, 15) = 0.14, *P =* 0.714; no group × day interaction, *F*(2, 30) = 1.77, *P =* 0.188. ***E***. Percentage of time spent in the reward-context during Pre-train and Post-train preference tests. Mixed ANOVA, group × test interaction, *F*(1, 15) = 19.41, *P =* 0.005. Sidak test for multiple comparisons, Post-train: *t(*30) = 5.19, *P* < 0.0001; Pre-train: *t*(30) = 0.5, *P =* 0.855; EYFP: *t*(15) = 3.86, *P =* 0.003; ArchT: *t*(15) = 2.33, *P=* 0.068. ***F***. Discrimination index of time spent in the reward-context. Unpaired t-test, *t*(15)= 3.74, *P =* 0.002. EYFP: *N* = 8 mice, ArchT: *N* = 9 mice. Data are presented as mean ± SEM.

These mice were then tested in a modified sucrose CPP paradigm^21,42^, conducted in an apparatus with two compartments (contexts) defined by unique visual and tactile cues. One day prior to the beginning of training, baseline exploration preference was determined in a 5-minute test (Pre-train test). Subsequently, on three daily training sessions, sucrose solution (10%) was placed in the context for which mice showed lower baseline preference and water was placed in the opposing context (Fig. 1B). Mice freely traversed between the reward- and neutral-contexts for 15 minutes, and light was delivered through the implanted optic fibres in a closed-loop manner exclusively during occupancy of the reward-context (15 mW of 532 nm continuous laser light measured from the fibre tip).

We did not observe any differences between ArchT and EYFP mice in the amount of sucrose consumed (Fig. 1C), or in the percentage of time spent in the reward-context throughout training days (Fig. 1D). These results indicate that the vHPC-NAc circuit does not contribute to the intrinsic motivational properties of sucrose reward. Following the final training session 3, we tested preference for the reward-context in the absence of sucrose and light delivery as a measure of memory for the context-reward association (Post-train test). While no group differences were observed during the Pre-train test, during the Post-train test, ArchT mice spent less time in the reward-context compared to EYFP mice (Fig. 1E,F). Furthermore, whereas EYFP mice spent more time in the reward-context during the Post-relative to the Pre-train test, this was not observed among ArchT mice (Fig. 1E). These results suggest that vHPC-NAc terminal inhibition during learning impaired memory encoding of the reward-context, revealing the necessity of activity in the vHPC-NAc circuit for associating rewards with the concurrent spatial context.

### vHPC-NAc projecting neurons are innervated by local CCK interneurons

Since we found that inhibiting the activity of vHPC-NAc pyramidal neurons disrupted the formation of contextual reward memory, we hypothesized that endogenous activity suppression by local GABAergic interneurons regulates learning^43,44^. Neural representations in the hippocampus rely on balanced microcircuit interactions between excitatory pyramidal neurons and inhibitory GABAergic interneurons^30^. In particular, we tested the impact of CCK interneurons since they provide strong feed-forward inhibition onto pyramidal neurons^45–48^, and have been implicated in dopamine-dependent behaviours^27^, and contextual memory^38,49^.

To investigate whether CCK interneurons monosynaptically innervate vHPC-NAc projecting cells, we used trans-synaptic pseudotyped rabies tracing^50^ (Fig. 2A,B). The retrograde virus CAV containing Cre (CAV-Cre) was infused into the NAc while a cre-dependent AAV vector containing the TVA receptor and optimized glycoprotein (oG) (AAV2/8-hsyn-DIO-TVA-2A-EGFP-2A-oG) were infused into the vHPC. After 2 weeks, G-deleted envelope-A pseudotyped rabies (EnvA-RVdG-mCherry) was infused into the vHPC, and mice were sacrificed 6 days later. In the vHPC, we observed cells positive for both EGFP and mCherry which constituted the ‘starter ‘ cell population, and cells singly labelled with mCherry which were considered presynaptic input cells (Fig. 2A). Immunostaining for GABA and CCK-8 revealed a significant proportion of the presynaptic GABA interneurons in the ventral CA1 (vCA1) and subiculum were CCK interneurons (Fig. 2B).

**Figure 2.**
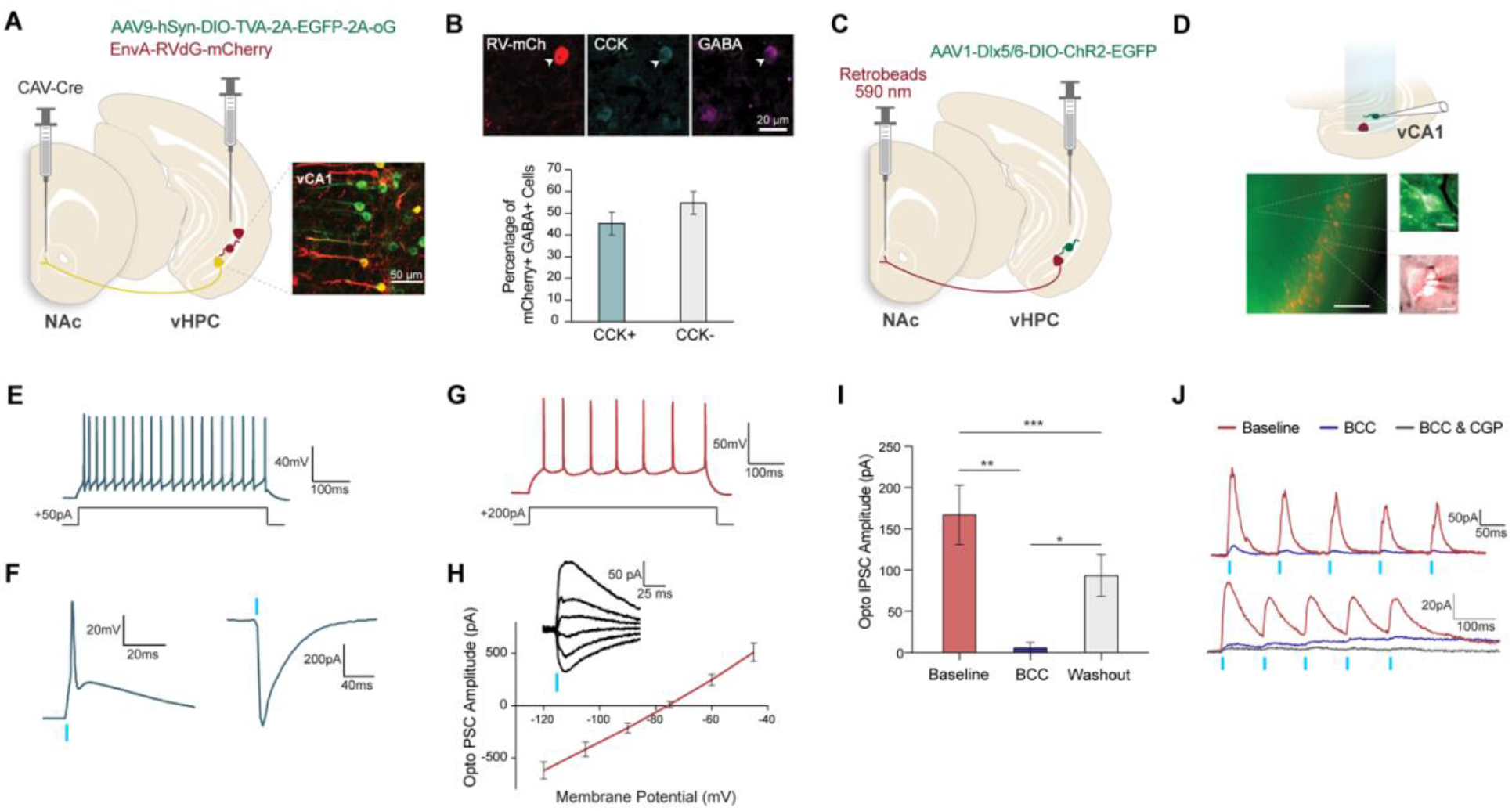
CCK interneurons in vHPC provide monosynaptic GABAergic inhibition of CA1 pyramidal neurons that project to nucleus accumbens (NAc). ***A***. Schematic of targeting approach for trans-synaptic pseudotyped rabies tracing from vHPC-NAc projecting neurons. Inset, representative image of starter cells double-labelled with EGFP and mCherry, and presynaptic input cells singly-labeled with mCherry in the vCA1. ***B***. Top, representative immunostaining for CCK-8 (cy5) and GABA (blue) against mCherry expression. Bottom, percentage of GABAergic presynaptic input cells (double-labelled with mCherry and GABA) that are immunoreactive for CCK-8. N = 4 mice. ***C***. Schematic of experimental strategy to express ChR2 in vHPC CCK interneurons and simultaneous retrograde tracing of vHPC-NAc projecting neurons with retrobeads. ***D***. Top, schematic of ex vivo electrophysiology in vCA1 with blue light illumination to visualize ChR2-EGFP+ CCK interneurons. Bottom, representative merged fluorescence and brightfield images of vCA1 (scale bar, 100 μ m). Top inset, ChR2-EGFP+ CCK interneuron (green, scale bar, 5 μ m). Bottom inset, merged fluorescence and IR-DIC images of a recording pipette patching a retrobead^vHPC→NAc^ pyramidal neuron (scale bar, 5 μ m). ***E***. Electrophysiological signature of a ChR2-EGFP+ CCK interneuron in response to a step of depolarizing current. *N =* 4 neurons from 4 mice. ***F***. Brief optogenetic stimulation (1 ms pulse, vertical blue line) excites a ChR2-EGFP+ CCK interneuron, in current-clamp from rest (left), and in voltage-clamp at −75 mV (right). ***G***. Electrophysiological signature of a retrobead^Vhpc→NAc^ vCA1 pyramidal neuron in response to a step pulse of depolarizing current. *N =* 16 neurons from 4 mice. ***H***. The optogenetically-evoked postsynaptic response reverses near the predicted equilibrium potential for a chloride ion current. Inset, example traces across a range of pyramidal cell holding potentials. ***I***. Pharmacological characterization confirms that optogenetically-elicited outward currents (Vh = − 60 mV) are suppressed by the competitive antagonist of GABA-A receptors (bicuculline: BCC, 3 μM). One-way repeated measures ANOVA, main effect of treatment, *F*(1.82, 7.26) = 30.64, *P* = 0.0003. Tukey ‘s multiple comparison tests: baseline vs. bicuculline: *P* = 0.0036; bicuculline vs. washout: *P* = 0.03. ***J***. Examples from two retrobead-positive vCA1 pyramidal neurons illustrating currents elicited by brief trains of optogenetic stimulation at baseline and upon blockade of GABA receptors.

To determine if CCK interneurons form functional connections with vHPC-NAc projecting cells, we performed ex vivo whole cell electrophysiological recordings from NAc projecting pyramidal neurons in the vCA1 while optogenetically stimulating CCK interneurons. To target the excitatory opsin channelrhodopsin (ChR2) to CCK interneurons, CCK-Cre mice were infused with an AAV vector containing Cre-dependent ChR2 driven by the GABA-specific Dlx5/6 promoter (AAV2/1-Dlx5/6-DIO-ChR2-EGFP) into the vHPC (Fig. 2C)^51^. To visualize vHPC-NAc projecting neurons in the same mice, red retrobeads were infused into the NAc. Blue light delivered to vHPC brain slices allowed ChR2-EGFP+ CCK interneurons to be visualized, patched, and stimulated (Fig. 2D-F). Their intrinsic properties included: input resistance 304 ± 60 MΩ, resting membrane potential −78 ± 5 mV, and spike amplitude 73 ± 3 mV (Fig. 2E). Stimulation of these ChR2-EGFP+ neurons with a short light pulse (1 ms) evoked an action potential under current clamp (Fig. 2F, left), and elicited a large inward current under voltage-clamp near resting membrane potential (− 911 ± 86 pA) (Fig. 2F, right).

Retrobead^vHPC→NAc^ neurons were distributed throughout the ventral CA1 and subiculum and were morphologically characteristic of pyramidal neurons (Fig. 2D). The intrinsic properties of retrobead^vHPC→NAc^ vCA1 pyramidal neurons included: input resistance 78 ± 6 MΩ, resting membrane potential −78 ± 1 mV, and spike amplitude 78 ± 4 mV (Fig. 2G). In response to optogenetic stimulation of the local axonal projections of ChR2-EGFP+ CCK interneurons (5 ms pulse), retrobead^vHPC→NAc^ neurons exhibited a postsynaptic current which reversed close to the predicted equilibrium potential for a chloride ion current (∼75 mV) (Fig. 2H). This current was strongly suppressed by the competitive GABA-A receptor antagonist bicuculline (3 μ M, 5 min) (Fig. 2I,J), a suppression that partially reversed after a brief washout period (< 10 min) (Fig. 2I). In a minority of neurons (2/7), a bicuculline-resistant outward current was recruited upon repeated stimulation (5 pulses, 10 Hz), which was sensitive to suppression by the GABA-B receptor antagonist CGP 35348 (1 μM) (Fig. 2J). Taken together, these findings suggest CCK interneurons provide functional GABAergic innervation of vHPC pyramidal neurons that project to the NAc.

### Intersectional genetic targeting of vHPC CCK Interneurons

To evaluate the contribution of CCK interneurons to behaviour, we expressed ArchT selectively in CCK interneurons to silence their activity in vivo. Due to nonspecific CCK neuropeptide expression in both GABA interneurons and pyramidal neurons^52^, CCK interneurons cannot be selectively targeted with conventional single recombinase methods. Therefore, we applied a dual recombinase intersectional approach^53,54^, using both the Cre/lox and Flpe/FRT systems, to genetically access CCK interneurons^37^. ArchT expression was targeted to CCK interneurons by crossing the CCK-Cre line with Dlx5/6-Flpe, and their double transgenic offspring with the Cre and Flpe-dependent RC::PFArchT-EGFP line. In the resulting CCK-Dlx5/6-FrePe triple transgenic mice (referred to as CCK-ArchT mice from here on), Flpe- and Cre-mediated excisions of the two transcriptional stop cassettes, the first flanked by loxP sites and the second by FRT sites, resulted in ArchT-EGFP expression selectively in CCK interneurons (Fig. 3A,B). Consistent with previous findings on the distribution of CCK interneurons in the hippocampus^55–57^, the labelled ArchT-EGFP+ neurons were observed across all layers of the ventral CA1 but were most abundant in the stratum pyramidale (Fig. 3C). A large majority of these neurons were immunoreactive for CCK-8 and GABA, while only a small percentage were immunoreactive for PV, SOM, or VIP (Fig. 3D), indicating high selectivity of the targeting approach for CCK interneurons.

**Figure 3.**
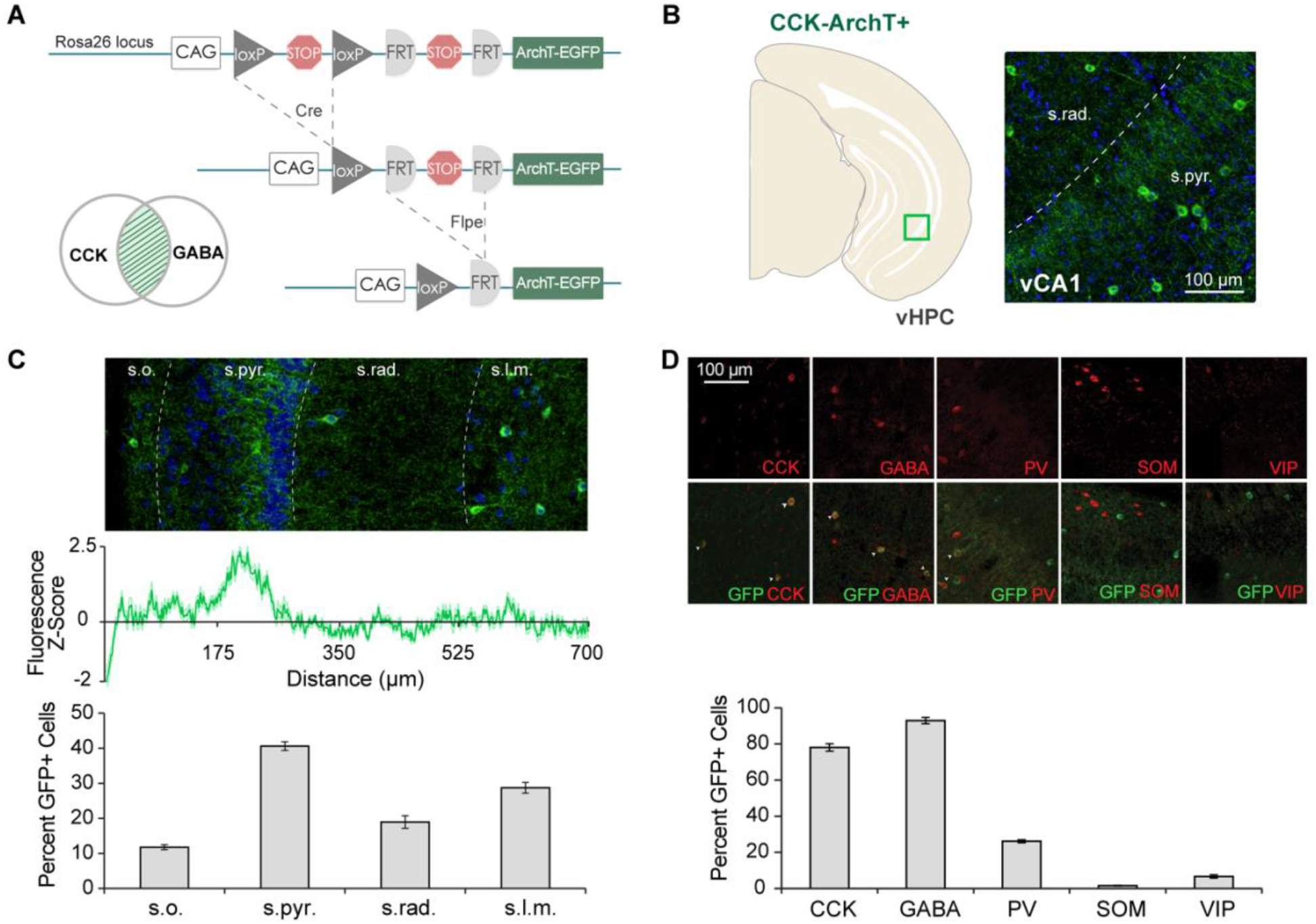
Intersectional genetic expression of ArchT in vHPC CCK interneurons. ***A***. Dual recombinase-responsive reporter allele, RC::PFArchT-EGFP, contains two transcriptional stop cassettes flanked by loxP and FRT sites. Cre- and Flpe-mediated excisions result in ArchT-EGFP expression in CCK interneurons. ***B***. Right, representative ArchT-EGFP expression (green) in vCA1 (boxed area in schematic) of a CCK-ArchT mouse. ***C***. Top, representative ArchT-EGFP expression across vCA1 strata. Middle, normalized fluorescence intensity (z-score) of ArchT-EGFP expression as a function of distance from the alveus. Bottom, percentage of ArchT-EGFP positive cells in each stratum of vCA1. One-way repeated measures ANOVA, *F*(1.31, 5.25) = 17.54, *P =* 0.0063, *N =* 5 mice. s.o. stratum oriens, s.pyr. stratum pyramidale, s.rad. stratum radiatum, s.l.m. stratum lacunosum-moleculare. ***D***. Top, representative immunostaining for GABA interneuron markers (red) against EGFP expression in CCK-ArchT mice. Bottom, percentage of ArchT-EGFP positive cells immunoreactive for given GABA interneuron marker. One-way ANOVA, *F*(4, 10) = 932.6, *P* < 0.0001, *N =* 3 mice. Data are presented as mean ± SEM.

## vHPC CCK interneurons regulate activity of vHPC-NAc projecting neurons in vivo

Next, we employed this genetic strategy to evaluate the effect of inhibiting vHPC CCK interneurons on the activity of vHPC-NAc projecting pyramidal neurons as mice explored a context containing reward. In CCK-ArchT mice, the retrograde tracer cholera toxin subunit B (CTB) conjugated to Alexa Fluor 555 was infused into the NAc shell resulting in CTB labeling of cell bodies in the ventral CA1 and subiculum (Fig. 4A,B,F). These mice were also implanted with bilateral optic fibres positioned above the vHPC (vCA1/subiculum regions) (Fig. 4A-C). First, mice were exposed to a context containing sucrose solution for 10 minutes during which light was delivered to inhibit vHPC CCK interneurons. Since we hypothesized that silencing CCK interneurons would disinhibit vHPC-NAc projecting neurons recruited by reward, a low concentration of sucrose solution (1%) was provided^58^. After context-reward exposure, mice were placed in their home cage for 60 minutes to allow cFos protein expression to accumulate and were then sacrificed^59^ (Fig. 4C). Inhibiting CCK interneurons in a reward-containing context significantly increased the density of cFos-positive neurons in the vHPC (Fig. 4D,E). Quantification of neurons that were double-labeled for CTB^vHPC→NAc^ and cFos revealed an increase in the percentage of CTB^vHPC→NAc^ neurons expressing cFos in CCK-ArchT+ compared to CCK- ArchT- mice (Fig. 4F,G). Therefore, inhibiting CCK interneurons in a context containing sucrose reward increased cFos expression in vHPC neurons that project to the NAc.

**Figure 4.**
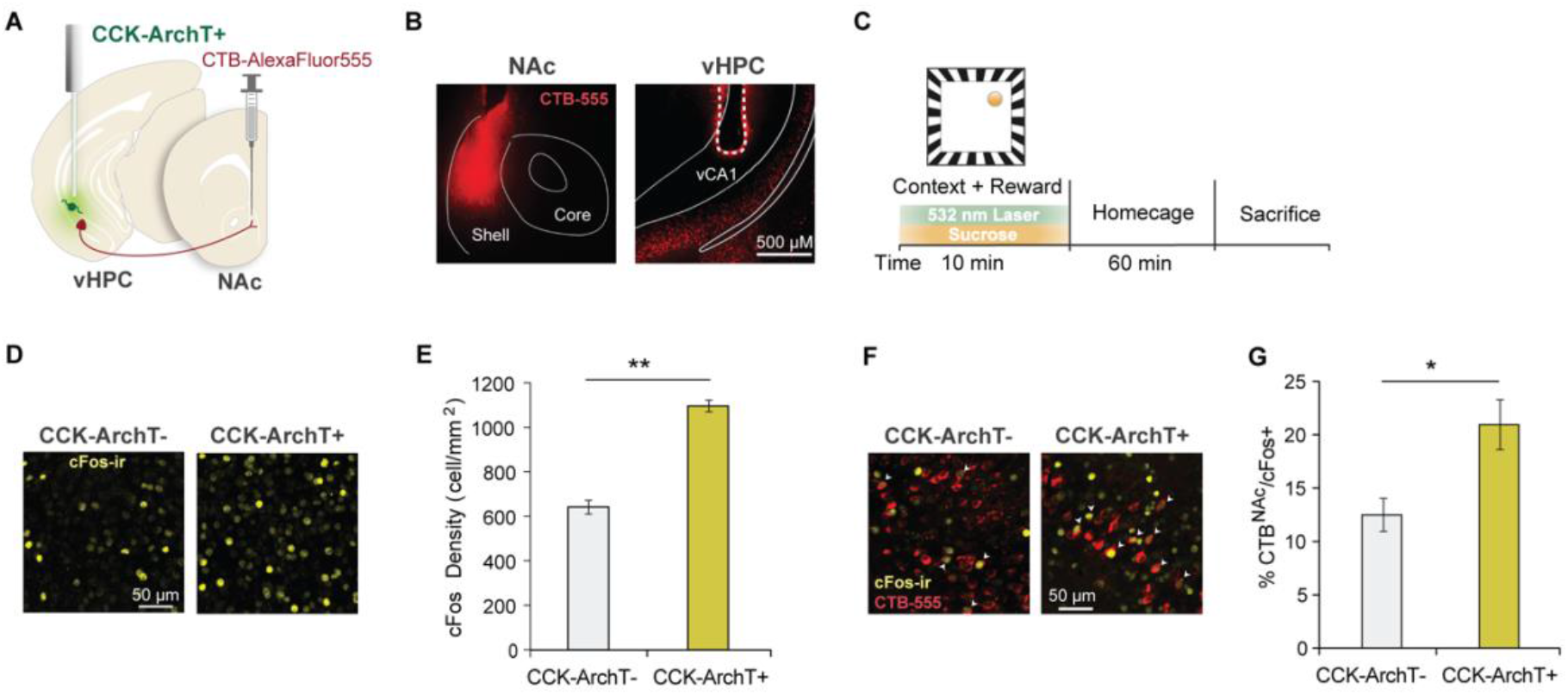
vHPC cFos expression following CCK interneuron inhibition during exposure to a context containing reward. ***A***. Schematic of retrograde tracing from nucleus accumbens combined with optogenetic inhibition of CCK interneurons in vHPC. ***B***. Left, representative image of CTB-Alexa-Fluor-555 infusion in NAc shell. Right, CTB ^vHPC→NAc^ retrogradely labeled cell bodies and optic fibre in vHPC. ***C***. Time course for optogenetic cFos experiment. Light was delivered during context and sucrose exposure. ***D***. Representative images of cFos-immunoreactivity in the vHPC from CCK-ArchT- (top) and CCK-ArchT+ (bottom) mice. ***E***. Density of cFos-immunoreactive cells in the vHPC. Unpaired t-test, *t*(9) = 4.28, *P =* 0.002; CCK-ArchT-: *N =* 6, CCK-ArchT+: *N =* 5. ***F***. Representative overlap between cFos-immunoreactivity and CTB^vH→NAc^ labeling in the vHPC of CCK-ArchT- (top) and CCK-ArchT+ (bottom) mice. ***G***. Percentage of CTB^vHPC→NAc^ cells in the vHPC immunoreactive for cFos. Unpaired t-test, *t*(8) = 3.03, *P =* 0.016; CCK-ArchT-: N = 5, CCK-ArchT+: N = 5. Data are presented as mean ± SEM.

### CCK interneuron inhibition enhances contextual reward memory

Since CCK interneurons were found to influence the activity of vHPC-NAc projecting neurons, we asked whether CCK interneurons may also be involved in the acquisition of contextual reward memory. We performed the sucrose CPP task as described in the previous section using 1% sucrose solution. Light was delivered to the vHPC of CCK-ArchT mice throughout the three training days exclusively during occupancy of the sucrose-containing context (Fig. 5A-C). During training, we did not observe any differences between CCK-ArchT+ and CCK-ArchT-mice in the amount of sucrose consumed (Fig. 5B), or in the percentage of time spent in the reward-context (Fig. 5C). In the place preference test, both groups spent a higher percentage of time in the reward-context during the Post-train test relative to the Pre-train test (Fig. 5D). However, CCK-ArchT+ mice exhibited significantly greater preference for the reward-context compared to CCK-ArchT-mice (Fig. 5D,E). To determine the specificity of this enhanced place preference memory, we subdivided the reward-context into quadrants and measured the time spent in each zone (Fig. 5F,G). Notably, CCK-ArchT+ mice spent a significantly greater percentage of time in the zone where the sucrose dish was previously located compared to CCK-ArchT-mice (Fig. 5G,H). These findings indicate that CCK interneuron inhibition strengthened learning of the context-reward association, resulting in enhanced memory of the location where reward was previously encountered.

**Figure 5.**
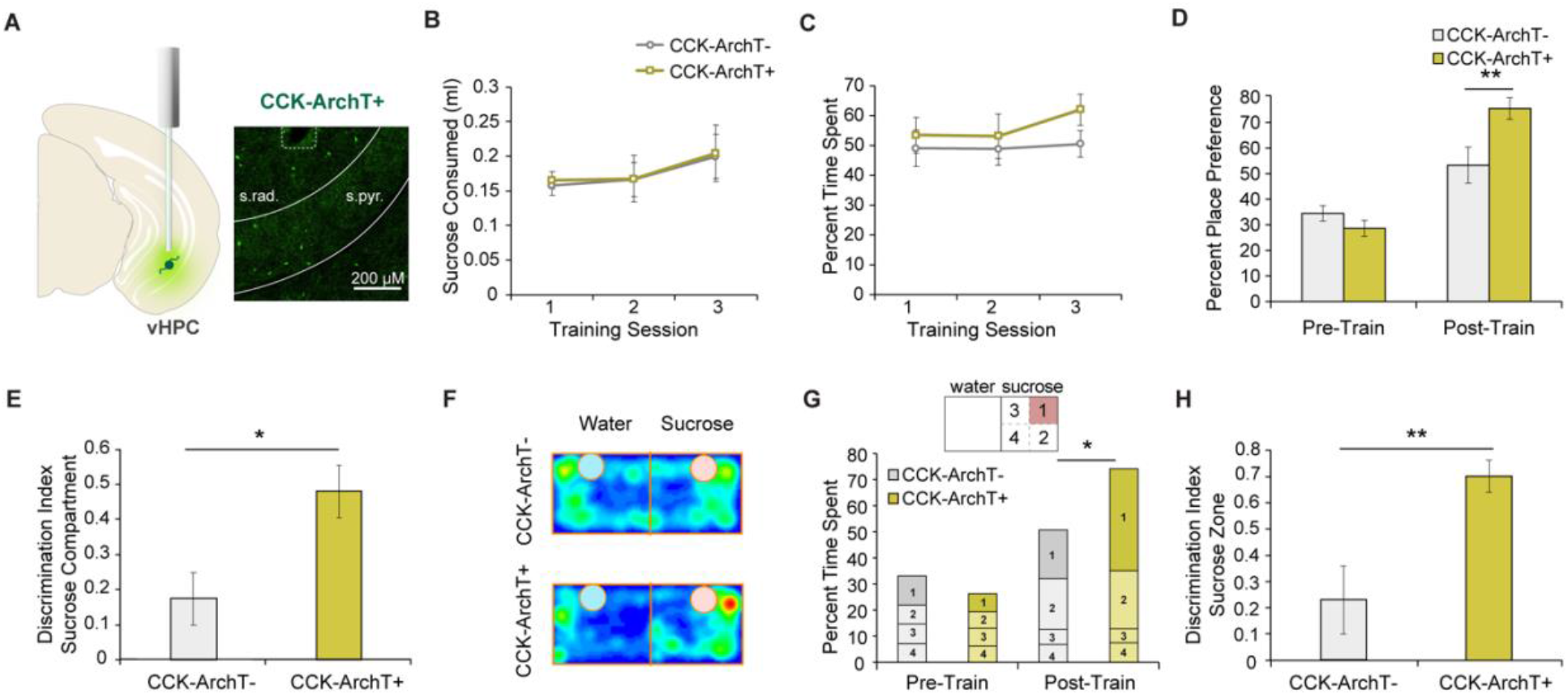
vHPC CCK interneuron inhibition during sucrose conditioned place preference training enhances memory at test. ***A***. Left, schematic of optogenetic targeting strategy of vHPC CCK interneurons. Right, magnified coronal image of vCA1 CCK interneurons expressing ArchT-EGFP. ***B***. Volume of sucrose solution consumed across training sessions. Mixed ANOVA, no main effect of group *F*(1, 15) = 0.03, *P =* 0.875, no group × day interaction, *F*(2, 30) = 0.01, *P =* 0.991. ***C***. Percentage of time spent in the light/sucrose-paired context across training sessions. No main effect of group, *F*(1, 15) = 0.84, *P =* 0.374; no group × day interaction, *F*(2, 30) = 0.68, *P =* 0.517. ***D***. Percentage of time spent in the reward-context during the Pre-train and Post-train preference tests. Mixed ANOVA, main effect of test, *F*(1, 15) = 46.05, *P* < 0.0001; group × test interaction, *F*(1, 15) = 10.34, *P =* 0.006. Sidak test, Post-train: *t*(30) = 3.59, *P =* 0.002, Pre-train: *t*(30) = 0.91, *P =* 0.6. ***E***. Discrimination index of time spent in the reward-context. Unpaired t-test, *t*(15)= 2.88, *P =* 0.011. ***F***. Group averaged heat maps of occupation time in the CPP apparatus during the Post-train test. ***G***. Percentage of time spent in individual zones of the reward-context during the Pre- and Post-train tests. Mixed ANOVA, group × context × zone interaction, *F*(3, 48) = 2.90, *P =* 0.044. Post-train: Sidak test, *t*(14.95) = 2.97, *P =* 0.038. ***H***. Discrimination index of time spent in the reward zone. Unpaired t-test, *t*(15) = 3.45, *P =* 0.004. CCK-ArchT-, N = 8; CCK-ArchT+, N = 9. Data are presented as mean ± SEM.

Alternatively, silencing CCK interneurons may have influenced preference behaviour during the Post-train test session by altering the reinforcing properties of the context itself (Britt et al., 2012), although, no direct effects of light delivery on preference were observed during sucrose CPP training. Another possibility is that mice may be detecting a mismatch in light-delivery during training and test that drives context exploration. We addressed these possibilities by inhibiting CCK interneurons in the same manner as the previous experiment but in the absence of sucrose during training. No significant differences were observed between CCK-ArchT+ and CCK-ArchT-mice in the percentage of time spent in the light-paired context during training (Fig. 6A,B), or during the Pre- and Post-train preference tests (Fig. 6C). Thus, CCK interneurons do not contribute to intrinsic preference behaviours in neutral contexts, and rather, influences memory-guided preference of contexts associated with reward.

**Figure 6.**
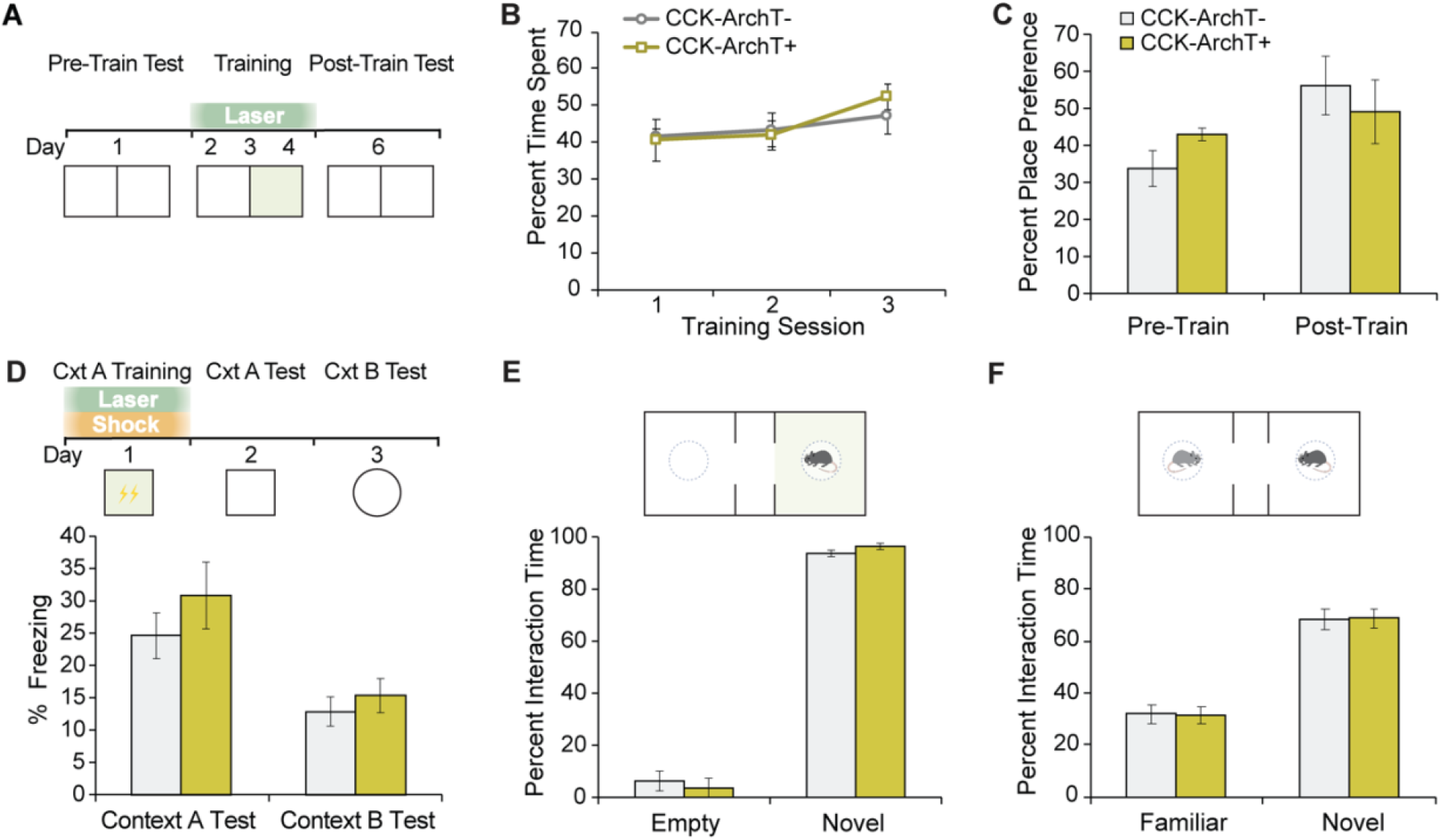
vHPC CCK interneuron silencing does not affect encoding of alternate forms of memory. ***A***. Behavioral paradigm for repeated real-time place preference with closed-loop optogenetic inhibition during training. ***B***. Percentage of time spent in the light-paired side across training sessions. Mixed ANOVA, no main effect of group, *F*(1,11) = 0.04, *P* = 0.842; no group × training day interaction, *F*(2,22) = 0.49, *P* = 0.62; CCK-ArchT+: *N =* 5 mice, CCK-ArchT-: *N =* 8 mice. ***C***. Percentage of time spent in the light paired side during the pre-training and post-training preference tests. Mixed ANOVA, no main effect of group, *F*(1,22) = 3.07, *P* = 0.094; no group × test day interaction, *F*(1,22) = 0.77, *P* = 0.391. ***D***. Behavioral paradigm for contextual fear conditioning with optogenetic inhibition during fear training (*Top*). Percentage of freezing in contexts A and B at test. Unpaired t-test, Context A: *t*(15)= 0.97, *P =* 0.346; Context B: t-test, *t*(15)= 0.70, *P =* 0.496. CCK-ArchT+: *N =* 8 mice, CCK-ArchT-: *N =* 9 mice. ***E***. Schematic of social memory conditioning with closed-loop optogenetic inhibition in the compartment containing the conspecific (*Top*). Percentage of time spent interacting with the empty cage or conspecific (*Bottom*). Mixed ANOVA, no main effect of group, *F*(1,16) = 0, *P >0*.*999*; no group × stimulus interaction, *F*(1,16) = 0.802, *P =* 0.384; CCK-ArchT+: *N =* 4 mice, CCK-ArchT-: *N =* 6 mice. ***F***. Schematic of the social recognition memory test (*Top*). Percentage of time spent interacting with the familiar or novel conspecific (*Bottom*). Mixed ANOVA, no main effect of group, *F*(1,16) = 0, *P >0*.*999*; no group × stimulus interaction, *F*(1,16) = 0.019, *P =* 0.893. Data are presented as mean ± SEM.

### CCK interneuron inhibition does not alter contextual fear or social memory

To test whether CCK interneuron inhibition of vHPC-NAc projecting neurons is specific for context-reward memory, we additionally investigated two other forms of memory. The vHPC has also been implicated in fear memory and social memory^4,13,20,60^. We evaluated whether vHPC CCK interneurons may be involved in the formation of contextual fear memory by optogenetically inhibiting CCK interneurons during contextual fear conditioning. No significant differences in freezing levels were found between CCK-ArchT+ and CCK-ArchT-mice during the test sessions in the fear conditioned context (Context A) or in a novel context (Context B) (Fig. 6D), indicating CCK interneuron inhibition did not affect contextual fear memory learning or specificity. Furthermore, we tested the role of CCK interneurons in the acquisition of social memory in a three-chamber social recognition memory task^61^. Light was delivered during the encoding phase only when mice occupied the chamber containing a social conspecific stimulus. CCK interneuron inhibition did not affect social interaction during the encoding phase (Fig. 6E), or during the social recognition memory test (Fig. 6F). Together, these experiments demonstrate that CCK interneuron inhibition of vHPC-NAc projecting neurons selectively controls context-reward memory.

## DISCUSSION

To successfully predict the availability of future rewards, animals must rapidly form and store internal representations of reward-containing environments. Yet, adaptive behaviour requires learning to be proportional to the probability and motivational value of rewards^62^, with maladaptive learning associated with conditions such as addiction and depression^63,64^.

Our study demonstrates that context-reward memory encoding is governed by a ventral hippocampal circuit in which CCK interneurons regulate CA1 pyramidal neurons projecting to the NAc. This work demonstrates that the strength of learning can be controlled by the balance of excitation in this circuit. Specifically, we found that, within the vHPC, inhibitory CCK interneurons monosynaptically innervate and modulate the excitability of pyramidal neurons projecting to the NAc. In reward-containing environments, inhibiting CCK interneuron activity increased the number of activated vHPC-NAc neurons and enhanced contextual reward learning. Meanwhile, inhibiting activity in the vHPC-NAc pathway had the opposite effect of impairing learning.

### Context-reward circuit in perspective

Pyramidal neurons in the vCA1 are heterogeneous in that they express unique genetic markers^65,66^, and send outputs to a diverse array of downstream targets^17^. Projection-defined pyramidal neurons may represent functional populations and have been reported to exhibit task-selective recruitment^10,13,18,22^. In the current study, we found that vHPC projections to the NAc are essential for associating a natural food reward with spatial contextual information. This finding aligns with previous studies demonstrating the involvement of this pathway in conditioned place preference for social reward^21^, and drug reward^22^. During reward learning, environmental cues gain incentive salience and come to elicit goal-directed behaviours^67^. The ability of spatial cues to drive reward seeking relies on coordinated activity between the hippocampus and NAc^68-70^. Both the dorsal and ventral hippocampus exhibit reward context-specific firing patterns following conditioning with drug or food rewards^9,10,71-73^, which in turn likely drives the context selectivity of NAc medium spiny neurons^72,74,75^. Importantly, our finding of enhanced memory for contexts associated with reward resulted from inhibiting vHPC CCK interneurons exclusively in the reward-paired context. Therefore, it is possible that CCK interneuron inhibition promoted the reward-context specific activity of NAc-projecting pyramidal neurons in the vHPC, leading to improved context encoding and discrimination. Future studies could evaluate this hypothesis by simultaneously inhibiting CCK interneurons and recording the activity of vHPC-NAc neurons in a contextual reward learning task.

### Clinical relevance of mechanism, caveats, and future directions

We optogenetically inhibited vHPC CCK interneurons during memory tests that were previously shown to be dependent on the vHPC^13,20,21,22,60^. Inhibiting CCK interneurons enhanced the formation of contextual reward memory. In contrast, the same manipulation did not affect the formation of contextual fear memory or social memory, indicating CCK interneurons do not strongly regulate the circuits mediating these behaviours. Specifically, the vHPC sends prominent projections to the basolateral amygdala to support contextual fear memory^4,13^, and to the medial prefrontal cortex for social memory^60^. Outputs of the vHPC to the NAc have also been reported to contribute to the retrieval of social memory^20^, but this was observed in a social recognition memory task that differed from the one used in the current study^60^. The selective involvement of CCK interneurons in contextual reward memory may indicate a biased functional connectivity with vHPC-NAc pyramidal neurons^29,76^. This possibility would need to be examined by comparing the inhibitory impact of CCK interneurons on multiple circuit-defined pyramidal neurons in vivo. Alternatively, vHPC CCK interneurons may play a more essential role in memories that are acquired gradually across repeated experiences as opposed to rapidly formed associative or incidental memories.

How would the unique properties of CCK interneurons allow for this population to act as a flexible gate in selecting pyramidal cell ensembles underlying context-reward memory? As we observed in the current study, CCK interneurons are distributed across layers in the CA1 and according to their axon localization are primarily dendrite-targeting or basket cells^55–57,77^. While CCK interneurons express several neuromodulatory receptors that would maintain their activity during wakefulness^48,78^, most intriguingly, they possess a unique ‘off ‘ switch, expressing high amounts of the CB1 receptor relative to other GABA interneuron subtypes and pyramidal neurons^79–81^. Stimulation of this receptor by endocannabinoids permits suppression of inhibition and long-term depression^38,82–85^. This mechanism would aid the encoding of context-reward associations in the hippocampus by amplifying the gain of spatial representations coinciding with reward^86^. Accordingly, we observed that inhibiting CCK interneurons both increased the number of activated NAc-projecting cells and improved contextual reward memory, consistent with an active role for this inhibitory circuit in regulating recruitment of pyramidal neurons to memory ensembles. Our preclinical work underscores the likely importance of CCK interneurons in optimizing context-reward memories, raising questions about the neuropathological consequences of damage or disrupted signalling to these neurons^87–90^.

## Conclusions

In summary, we identified a local inhibitory microcircuit in the ventral hippocampus in which CCK interneurons modulate the excitability of nucleus accumbens projecting pyramidal neurons to specifically control the strength of contextual reward learning. Thus, CCK interneurons may play a critical role for ensuring that contextual reward learning accumulates with experience over time, allowing for flexible reward-seeking behaviour in dynamic and unpredictable environments.

## MATERIALS AND METHODS

### Animals

We applied a dual recombinase intersectional approach (Kim et al., 2009), using both the Cre/lox and Flpe/FRT systems, to genetically access CCK interneurons (Whissell et al., 2015). Triple transgenic *CCK-Cre;Dlx5/6-Flpe;RC::PFArchT-EGFP* mice (termed CCK-ArchT mice) were generated as follows: *Dlx5/Dlx6-FLPe* mice were crossed with homozygous CCK-ires-Cre mice to generate double transgenic *Dlx5/6-Flpe;CCK-Cre* mice, which were then crossed with *RC::PFArchT-EGFP* mice (obtained from the laboratory of Dr. Itaru Imayoshi). Surgery and behaviour were performed on 4 –6 month old male mice.

### Stereotaxic Surgery

Mice were anaesthetized with isoflurane and mounted onto a stereotaxic frame. For optogenetic inhibition of ventral CA1 terminals in the nucleus accumbens, AAV2/5-CaMKIIα-eArchT3.0-EYFP or AAV2/5-CAMKIIα-EYFP (0.3 μL, UNC Vector Core) was bilaterally infused into the ventral hippocampus (AP: -2.90 mm, DV: -4.90, ML: ± 3.30 mm), and optic fibres were implanted bilaterally above the nucleus accumbens shell AP: 1.60 mm, DV: -4.00, ML: ±0.60 mm). For transsynaptic rabies tracing experiments, wildtype C57BL6/J mice were unilaterally infused with CAV2-cre (0.5 μL, Plateforme de Vectorologie de Montpellier) into the NAc (AP: 1.60 mm, DV: -4.75, ML: ±0.60 mm), and with AAV2/8-EF1α-DIO-TC66T-2A-EGFP-2A-oG (0.3 μL, GT3 Viral Core Facility of the Salk Institute) into the left or right ventral CA1/subiculum. Two weeks later, EnvA-RVdG-mCherry (0.7 μL, GT3 Viral Core Facility of the Salk Institute) was infused into the same hemisphere of the ventral CA1/subiculum. For *ex vivo* electrophysiology experiments, CCK-Cre mice were bilaterally infused with AAV2/1-Dlx5/6-DIO-ChR2-EGFP (0.3 μL, Vigene Biosciences, custom production) into the ventral CA1/subiculum and with rhodamine retrobeads (590 nm, 1 μL, Lumafluor Inc.) into the nucleus accumbens. For optogenetic experiments with CCK-Cre/DLX5/6-Flp/PFArchT mice, optic fibres were implanted bilaterally above the ventral CA1/subiculum region (AP: -2.90 mm, DV: -4.10, ML: ± 3.30 mm). For retrograde tracing experiments in combination with cFos labeling, cholera toxin subunit B conjugated to Alexfluor-555 (0.7 μL, Molecular Probes) was infused unilaterally into the NAc of CCK-ArchT mice. Infusions were made via an internal cannula (31 gauge) in the target region connected by tubing to a 10 uL Hamilton syringe. The internal cannula was left in place for 10 minutes after infusion to prevent solution backflow. Optic fibres were secured to the skull using dental cement (RelyX Unicem; 3M). After surgery, mice were individually housed and allowed a minimum of 1 week to recover before behavioural experiments. The brain coordinates described are in reference to Paxinos and Franklin (2007).

### Behavioural Apparatuses and Testing Procedures

One week after recovery from stereotaxic surgery, mice were individually handled for 5 minutes on 2 consecutive days, and subsequently acclimated to being connected to the optical patch cords for 5 minutes in the behavioural testing room. Behavioural tests were conducted under room lighting (100 lux), and tracking and scoring were done using ANY-maze™ (Stoelting Co.) unless specified otherwise. For optogenetic inhibition, 532 nm of laser light was set to a power of 15 mW from the fibre tip and delivered continuously, either throughout the apparatus or in a particular compartment as described in the specific behavioural methods below. Apparatuses were cleaned with 70% ethanol before the introduction of each individual mouse, unless specified otherwise.

### Sucrose Conditioned Place Preference

To test for changes in learning of context-reward associations, sucrose conditioned place preference was performed. Conditioned place preference for sucrose solution (1% or 10% in water) was tested in a rectangular chamber comprised of two contexts (each 25 cm L × 20 cm W × 30 cm H) with mice having free access to both contexts. The walls of each context had a pattern that was either black and white horizontal stripes or black spots, and the floor was one of two metal textures. One day prior to training, mice were tested for their baseline preference of the two contexts in a 5-minute session (Pre-Train Test). During training, the less preferred context contained sucrose solution and the preferred side contained water. A petri dish (6 cm diameter) was filled with each solution (10 ml) and placed into either context at the far opposing walls. Mice were placed into the light-unpaired side facing the wall and allowed to explore for 15 minutes. Light was delivered upon entry into the sucrose-paired context as measured from the body ‘s centre point. Solutions were weighed before and after training to measure the amount of consumption during training. Mice were trained over 3 consecutive sessions. Forty-eight hours after the last session, a 5 minute preference test was conducted (Post-Train Test). For this test, mice were first food and water deprived for 16 hours.

### Contextual Fear Conditioning

The contextual fear task was conducted in arenas placed inside of sound-attenuating boxes (Med Associates Inc.). Two unique arenas (contexts) were used which differed in their visual, tactile, and olfactory cues. The fear conditioning context (context A) was a square arena (30 cm L × 24 cm W x× 21 cm H) that had walls constructed of metal and clear plastic, a metal grid floor (19 stainless steel rods), and a house light, and was cleaned with 70% ethanol. The neutral context (context B) was semicircular in shape and was made of smooth white plastic (30 cm wide) and was cleaned with 4% acetic acid. During conditioning on day 1, mice were placed into context A for a total of 300 seconds during which 2 foot-shocks (2 s, 0.7 mA) were delivered at timepoints 180 and 240 seconds into the session. The next day, mice were returned to context A for 300 seconds during which freezing was measured as an index of contextual fear memory. On day 3, mice were placed into the novel context B for 300 seconds during which freezing was measured as index of contextual fear memory generalization. Freezing (minimum bout duration of 0.5 s) was automatically scored using tracking software (FreezeFrame 4, Coulbourne Instruments).

### Social Recognition Memory

Social memory was tested in a three-chamber social recognition memory test (Moy et al., 2004). The apparatus was made of clear Plexiglas and consisted of three chambers, each measuring 40 cm L × 20 cm W × 40 cm H. The two side chambers contained a cylindrical wire cage (10.5 cm D × 11 cm H, 1 cm bar spacing). Subjects were able to freely move between chambers through an opening (5 cm × 40 cm) in the partitioning walls. Subject mice were placed into the centre chamber and allowed to acclimate to the entire apparatus for 10 minutes. Following acclimation, a 4-week old adolescent male C57BL/6 mouse, unfamiliar to the subject, was placed into a cage in one of the side chambers and interaction was permitted for 5 minutes. Immediately following this phase, the subject was confined to the centre while a second unfamiliar mouse was placed in the previously empty cage. Interaction was again permitted for 5 minutes to test for social recognition memory. Interaction was manually scored by key presses as any directed nose contact with the target mouse or cage. The initial unfamiliar mouse and chamber side were counterbalanced within control and CCK-ArchT groups.

### cFos Behavioural Procedures

To determine whether CCK interneuron inhibition during context-reward association learning affects vHPC neural activity, we analyzed cFos expression following exposure to a context containing reward with optogenetic inhibition of CCK interneurons. Mice were first individually handled for 5 minutes each on 3 consecutive days followed by a 2-hour habituation period to the behavioural testing room. On the fourth day, mice were connected to the optical patch cords and allowed to acclimate for 5 minutes in the home-cage. They were then habituated to the arena (25 cm L × 20 cm W x× 30 cm H) without reward for 5 minutes and returned to the home-cage in the testing room for another 2-hour period. The next day, mice were habituated to the testing room for 2 hours and were then placed into the context with 1% sucrose reward and light delivery for 10 minutes. After testing, mice were returned to their home-cage and sacrificed 60 minutes later.

### *Ex vivo* Electrophysiology

#### Slice preparation

Mice were sacrificed at postnatal day 90– 120 and 1– 2 weeks after the final stereotaxic surgery in order to prepare acute brain slices for optophysiological recording. We injected chloral hydrate solution (400 mg/kg) interperitoneally before decapitation. The brain was extracted into ice-cold oxygenated ACSF-sucrose solution (254 mM sucrose, 10 mM D-glucose, 24 mM NaHCO_3_, 2 mM CaCl_2_, 2 mM MgSO_4_, 3 mM KCl, and 1.25 mM NaH2PO4, pH 7.4). A Dosaka Linear slicer was used to prepare ventrohippocampal coronal slices (∼ 400 μ m thick, Bregma -2.9mm to -3.8mm); each slice was transferred to a chilled ACSF-sucrose solution filled petri dish and halved at the midline. Slices were then transferred to an oxygenated recovery chamber filled with 30°C ACSF solution (128 mM NaCl, 10 mM D-glucose, 26 mM NaHCO_3_, 2 mM CaCl_2_, 2 mM MgSO_4_, 3 mM KCl, and 1.25 mM NaH_2_, PO_4_,, pH 7.4). After >1 hour of post-slicing recovery, slices were placed on to a slice chamber and perfused with 30° C ACSF for optophysiological recordings.

### Electrophysiological recordings

EGFP-positive CCK interneurons and retrobead-positive CA1 pyramidal cells in ventrohippocampal brain slices were identified respectively using a white-light collimated LED and FITC and TRITC filter cubes. Labelled neurons were targeted for patching using IR-DIC via 60x lens on a BX51WI microscope (Olympus). Pipettes (∼2·4 M) contained potassium gluconate (120 mM K-gluconate, 5 mM KCl, 2 mM MgCl2, 4 mM K2-ATP, 0.4 mM Na2-GTP, 10 mM Na_2_-phosphocreatine, and 10 mM HEPES buffer; pH 7.3). Whole cell patch clamp recordings were conducted using a HEKA EPC10 amplifier and Patchmaster software and were analyzed using Clampfit-10.7 software. Channelrhodopsin was stimulated with light via a 445/45 nm excitation filter, delivered in 1 ms pulses to stimulate the somata of ChR2-EGFP+ CCK interneurons or 5 ms pulses to excite their axons terminating on retrobead^vHPC→NAc^ vCA1 pyramidal neurons.

### Pharmacology

Drug manipulations to the slice during optophysiological experiments were administered via ACSF perfusion. Antagonists included the GABA-A blocker bicuculline methiodide (BCC, 3 μM, Tocris) by itself or, in a subset of neurons, BCC in combination with the GABA-B blocker CGP 35348 (CGP, 1 μM; Tocris). Antagonist was pre-applied for 5 minutes before and co-applied during the relevant optophysiological recording. To examine the recovery of the optophysiological effect upon partial washout, additional recordings were performed in a subset of neurons ∼5-10 mins after BCC application ended.

### Histology

Mice were transcardially perfused with PBS, pH 7.4, followed by 4% PFA. Brains were extracted and postfixed overnight in 4% PFA at 4°C and then cryoprotected with PBS containing 30% sucrose. Brains were sectioned coronally at 40 μm thickness using a cryostat (Leica Microsystems, CM 1520). For immunostaining, free-floating brain sections were blocked with 5% normal donkey serum in 0.1% Triton X-100 in PBS (PBS-T) for 2 h. Sections were then incubated for 48–72 h at 4°C with PBS-T containing a combination of the following primary antibodies: rabbit polyclonal anti-CCK (1:1000, Sigma Millipore, C2581; Frontier Institute, Af350), rabbit polyclonal anti-GABA (1:1000, Sigma Millipore, A2052), rabbit polyclonal anti-PV antibody (1:1000, Abcam, ab11427), rabbit polyclonal anti-vasoactive intestinal peptide (VIP) (1:500, Immunostar, 20077), rat polyclonal anti-somatostatin (1:500, Millipore, MAB354), rabbit polyclonal anti-cFos (1:1000, Santa Cruz Biotechnology), chicken polyclonal anti-GFP (1:1000, Abcam, ab13970), and goat polyclonal anti-mCherry (1:1000, Sicgen, AB0040-200). Primary antibody incubation was followed by incubation for 2 h at room temperature with PBS-T containing the following secondary antibodies: AlexaFluor-488-conjugated donkey anti-rabbit (1:1000, Jackson ImmunoResearch Laboratories, 711545152), AlexaFluor-594-conjugated donkey anti-rabbit (1:1000, Jackson ImmunoResearch Laboratories, 715515152), Cy5-conjugated donkey anti-rabbit (1:1000, Jackson ImmunoResearch Laboratories, 711175152), DyLight-405-conjugated donkey anti-rabbit (1:500, Jackson ImmunoResearch Laboratories, 711475152), AlexaFluor-488-conjugated donkey anti-chicken (1:1000, Jackson ImmunoResearch Laboratories, 703545145), and AlexaFluor-594-conjugated donkey anti-goat (1:1000, Jackson ImmunoResearch Laboratories, 705515147). For cell counting experiments, every fourth section of the vHPC was collected and immunostained. The sections were mounted and imaged on a confocal laser scanning microscope with a 20× objective (Leica LSM 800). Fluorescent cells were counted in the ventral CA1 and subiculum (bregma -3.30 mm to -3.8 mm) in a 300 × 700 μM area.

### Statistical Analysis

Data were analyzed using One-way ANOVA, repeated measures ANOVA, Mixed ANOVA, and paired and unpaired Student ‘s t-tests. Where appropriate, ANOVAs were followed by planned pairwise comparisons. Statistical analyses were performed using IBM SPSS Statistics Version 21 and GraphPad Prism 6.

